# Loss of the homologous recombination gene *rad51* leads to Fanconi anemia-like symptoms in zebrafish

**DOI:** 10.1101/095646

**Authors:** Jan Gregor Botthof, Ewa Bielczyk-Maczyńska, Lauren Ferreira, Ana Cvejic

## Abstract

*RAD51* is an indispensable homologous recombination protein, necessary for strand invasion and crossing over. It has recently been designated as a Fanconi anemia (FA) gene, following the discovery of two patients carrying dominant negative mutations. FA is a hereditary DNA repair disorder characterized by various congenital abnormalities, progressive bone marrow failure and cancer predisposition. In this paper, we describe the first viable vertebrate model of *RAD51* loss. Zebrafish *rad51* loss-of-function mutants developed key features of FA, including hypocellular kidney marrow, sensitivity to crosslinking agents and decreased size. We show that some of these symptoms stem from both decreased proliferation and increased apoptosis of embryonic hematopoietic stem and progenitor cells. Co-mutation of *p53* was able to rescue the hematopoietic defects seen in the single mutants, but led to tumor development. We further demonstrate that prolonged inflammatory stress can exacerbate the hematological impairment, leading to an additional decrease in kidney marrow cell numbers. These findings strengthen the assignment of *RAD51* as a Fanconi gene and provide more evidence for the notion that aberrant p53 signaling during embryogenesis leads to the hematological defects seen later in life in FA. Further research on this novel zebrafish FA model will lead to a deeper understanding of the molecular basis of bone marrow failure in FA and the cellular role of RAD51.

**Significance statement:** The homologous recombination protein RAD51 has been extensively studied in prokaryotes and lower eukaryotes. However, there is a significant lack of knowledge of the role of this protein and its regulation in an *in-vivo* context in vertebrates. Here we report the first viable vertebrate mutant model of *rad51* in zebrafish. These mutant fish enabled us to confirm for the first time the recently discovered role of *RAD51* in Fanconi anemia pathogenesis. We report that p53 linked embryonic stem cell defects directly lead to hematological impairments later in life. Co-mutation of *rad51* with *p53* rescues the observed hematological defects, but predisposes the fish to early tumor development. The application of this model opens new possibilities to advance Fanconi anemia drug discovery.

## Introduction

Fanconi anemia (FA) is a hereditary DNA repair disorder characterized by various congenital abnormalities, progressive bone marrow failure (BMF) and cancer predisposition(1). It is caused by mutations in one of 21 genes in the FA pathway(2–4). The FA pathway has been shown to be the major route for the removal of interstrand crosslinks (ICL) - DNA lesions that prevent replication and transcription by inhibiting DNA strand separation(5, 6). When the pathway is defective, these structures cannot be removed, potentially leading to cell death(7). Indeed, sensitivity to crosslinking agents, such as mitomycin C, is an absolute diagnostic criterion of FA(8).

Although FA is characterized by remarkable phenotypic heterogeneity, FA patients usually succumb to the depletion of hematopoietic stem and progenitor cells (HSPCs) in their bone marrow, leading to pancytopenia and complete BMF. Therefore, bone marrow transplantation (BMT) is the only modality that offers a potential cure of hematopoietic defects but is itself associated with considerable morbidity(9, 10). Interestingly, a decrease in HSPCs (CD34+ cells) is already apparent in FA infants even before the first hematological symptoms appear(11). This led to the hypothesis that FA originates from defects during the formation of the initial HSPC pool, presumably due to an overactive p53/p21 response and cell cycle arrest(11). In agreement with this, FA mice have considerably smaller fetal livers than their healthy siblings(12). It remains unclear however, at which stage during embryonic development these defects appear and how perturbation in the production of embryonic HSPCs relates to the phenotype seen in adulthood.

Due to the role FA genes play in the repair of interstrand crosslinks, DNA damaging agents causing ICLs have been proposed as a major cause of BMF, with small aldehydes being the most likely candidates. Co-mutation of genes in the FA pathway and aldehyde metabolizing genes (*Aldh2* and *Adh5*) resulted in significant reduction of HSPCs and BMF in double mutant mice(13–17). In addition, FA patients lacking *ALDH2* show a more severe phenotype(18, 19). Apart from their hypersensitivity to crosslinking agents, FA cells also react excessively to pro-apoptotic cytokines such as IFNγ and TNFα(20–25). However, the role of cytokines in the etiology of BMF remains controversial(26–29).

In the last two years, a novel FA subtype associated with dominant-negative mutations in *RAD51* was reported, leading to the designation of *RAD51* as *FANCR(30–32)*. It has been shown to be involved in protecting broken down replication forks from excess processing by nucleases, linking the FA pathway with RAD51/BRCA2(30, 33). *In-vivo* studies of Rad51 have previously been very difficult, as mice lacking the protein invariably die during early embryogenesis(34, 35).

In this study, we characterized the first viable vertebrate model of Rad51 loss. Indeed, our zebrafish *rad51* loss of function mutant recapitulates many congenital and haematological features of FA. We provide the first *in-vivo* evidence that decreased HSPC numbers during embryonic development directly lead to the later bone marrow defects in FA. Finally, we show that *rad51* mutants do not overproduce inflammatory cytokines, but are more sensitive to them and that prolonged inflammatory stress can further reduce marrow cellularity.

## Results

### The *rad51^sa23805^* allele leads to complete loss of functional Rad51

To study the function of Rad51 in hematopoiesis, we obtained fish carrying the *rad51^sa23805^* allele from the Sanger Institute Zebrafish Mutation Project(36). The *rad51^sa23805^* allele has a C>T mutation at codon 203 in exon 7, which leads to a premature stop codon in the region of the AAA+ ATPase domain. In contrast to mice lacking Rad51, which invariably die during early development(34, 35), fish carrying homozygous copies of the *rad51^sa23805^* (referred to as *rad51*-/- for brevity in the text) survive to adulthood. However, all surviving adults undergo sex reversal and are infertile males, lacking mature spermatozoa in the testes (Figure S1A and B).

To ensure that the mutation is actually functional, we carried out a Western blot on testicular tissue, since Rad51 is highly expressed in germ cells in zebrafish as is in other species(37, 38). This confirmed that the full-length protein is lost in *rad51*-/- fish (Figure 1A).

**Figure 1.**
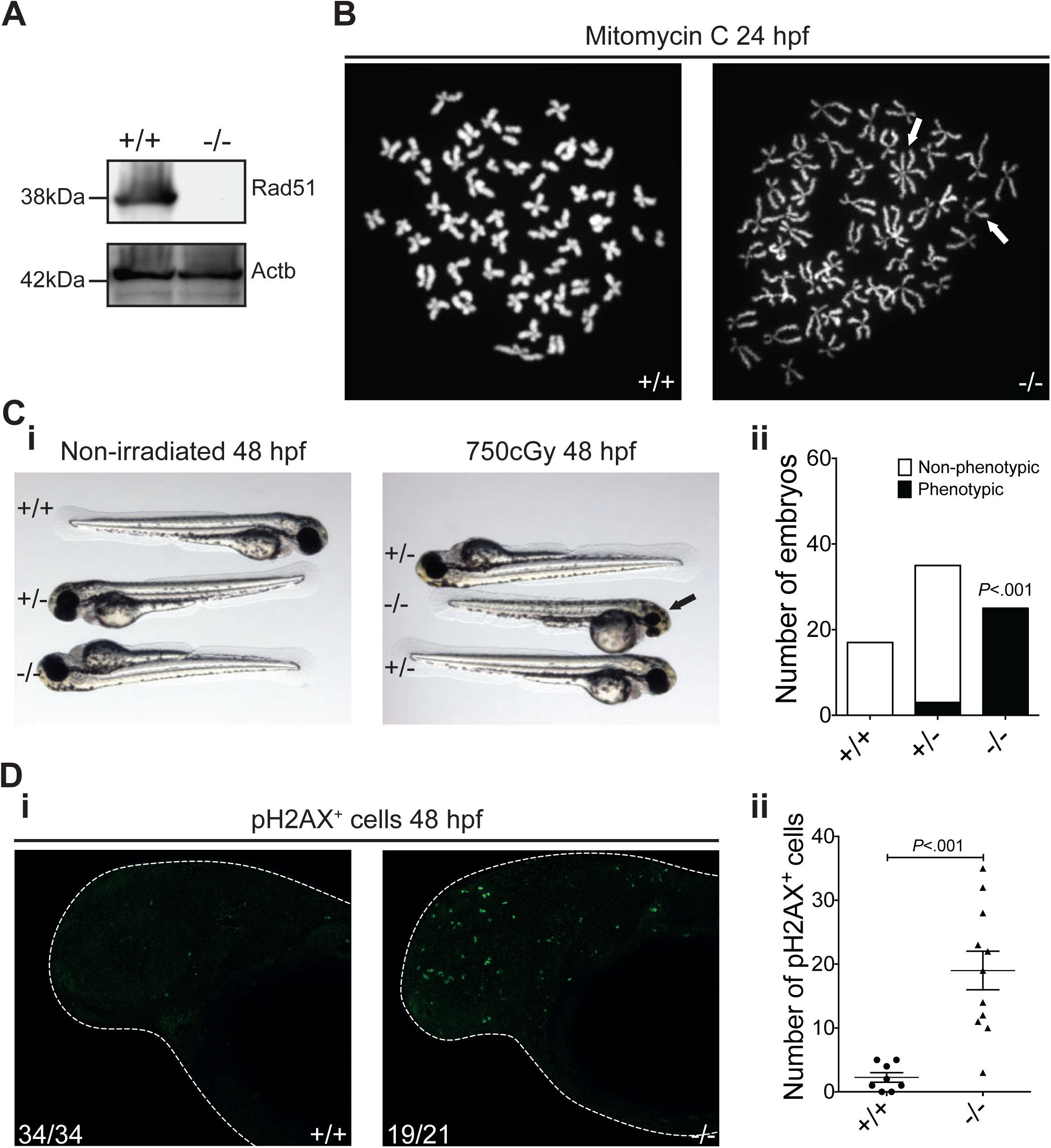
The *rad51^sa23805^* allele leads to loss of Rad51 protein and causes DNA damage sensitivity. (A) Western blot showing the expression of Rad51 in testes extracts of wild type (wt) and mutant zebrafish. Beta-actin was used as a loading control. (B) Chromosome spreads of 24 hpf wild type and mutant embryos treated with 1μg/ml mitomycin C for 20 hours. White arrows indicate characteristic damage (chromosome breaks and radial structures) in response to crosslinking agents. (C) Comparison of the response of 48 hpf wild type and mutant embryos to irradiation (i). The black arrow indicates the small head and eye phenotype, which is quantified in ii. Two-tailed Fisher’s exact test pooling wild types and heterozygotes as control group, *P*<0.001, n=67. (D) Immunostaining for pH2AX in wild type and mutant embryos with representative embryos (i) and quantification of foci (ii). White arrows indicate example foci. Two-tailed Student’s t-test, *P*<.0001, n_+/+_=8, n_−/−_=11.

### Lack of Rad51 in embryos leads to increased DNA damage sensitivity

Rad51 is essential for the repair of interstrand crosslinks (ICLs) and double stranded breaks via homologous recombination (HR). In ICL repair, it plays a role in both HR-linked processes HR, as well as HR-independent steps(30–32). To investigate the role of Rad51 in the repair of ICLs, we treated embryos resulting from an incross of *rad51*+/− parents with the crosslinking agent mitomycin C (MMC). After treatment, the tails of the embryos were used for chromosome spreads, whereas the heads were kept for genotyping. The chromosome spreads revealed that MMC treatment led to the formation of characteristic chromosomal abnormalities, including breaks and radial structures, exclusively in the embryos lacking Rad51 (Figure 1B).

We also examined the role of Rad51 in the repair of other forms of double stranded breaks not involving crosslinking agents. For this, we irradiated 24 hour old embryos from an incross of *rad51*+/− parents with gamma radiation and examined them at 48 hpf. The irradiated embryos were then blindly imaged, scored and genotyped. The *rad51*-/- embryos developed small eyes and heads in response to radiation (Figure 1 Ci). This phenotype was limited to the *rad51*-/- embryos (Figure 1 Cii), showing that they are more sensitive to irradiation than wt siblings. In addition, pH2AX (a marker of double stranded breaks) immunostaining of non-irradiated embryos revealed extensive DNA damage in embryos lacking Rad51 in comparison to their wt siblings (Figure 1Di and ii). Taken together, these data suggest that *rad51*-/- embryos are hypersensitive to DNA damage.

### Rad51 mutants recapitulate many congenital and hematological features of Fanconi anemia

*RAD51* mutations were recently linked to FA in humans in two case reports(30, 31). Therefore, we examined the *rad51* mutant fish for congenital and hematological features associated with FA.

A common congenital symptom of FA is decreased height(1). On account of that, we measured body length of fish with and without *rad51* mutation throughout development. Although embryonic development was not affected by the mutation, the size of *rad51*-/- fish was decreased compared to their wt siblings, starting from around 23 dpf (Figure S2A). The size reduction during the larval period was maintained to adulthood with *rad51* mutant fish being on average 10% shorter than their wt siblings (Figure S2B and C). In addition, *rad51*-/- embryos and larvae developed microphthalmia, another distinct feature of FA patients (Figure S3).

To assess whether loss of *rad51* affects adult hematopoiesis, we inspected the kidney marrow (WKM) of zebrafish, which is analogous to bone marrow in mammals. Unlike in mammals, there is only one kidney in zebrafish, which is the only site of adult haematopoiesis. Importantly, there is no space restriction of the marrow by the rigid bone, as kidney tissue is very spongy and flexible. We first obtained H&E stained histological sections of the kidney, which showed no noticeable qualitative differences between *rad51+/+* and -/- fish (Figure 2A). However, even though the morphology appeared normal, kidney size and cell number were considerably (~50%) decreased in *rad51*-/- fish (Figure 2Bi and ii). The kidney cellularity of *rad51*-/- fish gradually decreased with aging, but at the same rate as in wt fish (Figure 2Bii) suggesting that the mutants established steady state hematopoiesis despite displaying kidney hypocellularity. Although *rad51* mutants did not develop cytopenia in the peripheral blood (PB) (Figure 2C), they did accumulate macrocytic erythrocytes in the PB (Figure 2D), suggesting a gradual worsening of the phenotype over time.

**Figure 2.**
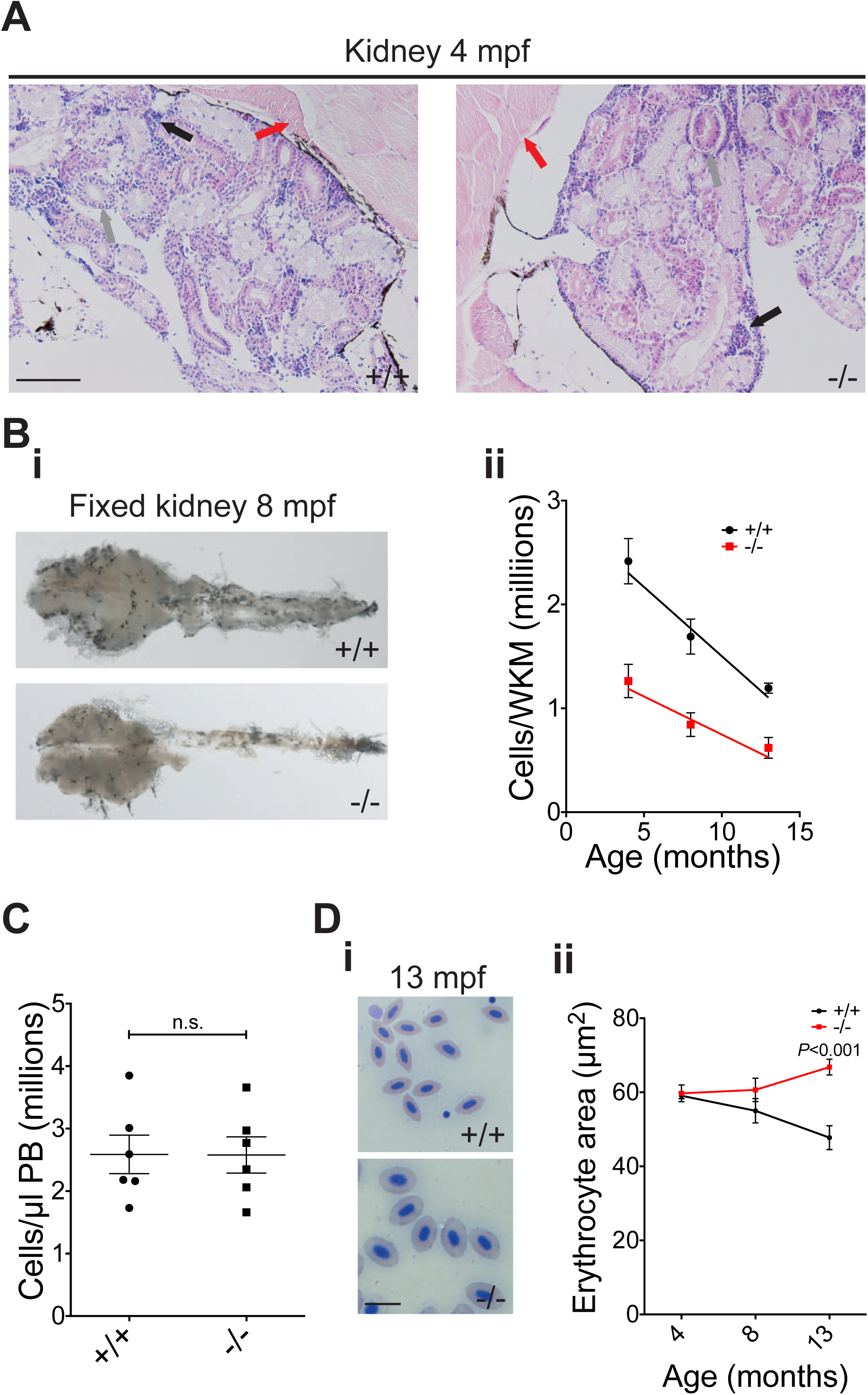
Adult *rad51* mutant fish display kidney marrow cytopenia. (A) H&E stained histological sections of 4 mpf wild type and mutant kidneys using a 20X objective. Muscle (red arrow), ducts/tubules (grey arrow) and hematopoietic kidney marrow (black arrow) can be seen. The scale bar is 100 μm. (B) Fixed 8 mpf wild type and mutant kidneys (i); quantification of the number of total cells per freshly isolated kidney at different ages using a hemocytometer (ii). Two-way ANOVA was used and type III model fit (Armitage 2001). The test shows a significant influence of age (F (1,50)=18.23, *P*<.0001) and mutation status (F (1,50)=10.87, *P*=.0018) on phenotype. 4 mpf n_+/+_=6, n_−/−_=6; 8 mpf n_+/+_=16, n_−/−_=16; 13 mpf n_+/+_=6, n_−/−_=4. (C) Quantification of peripheral blood (PB) cells in wild type and mutant fish at 4 mpf. Two-sided T-test, n_+/+_=6, n_−/−_=6. (D) In i, blood smears of 13 mpf wild type (top) and mutant fish (bottom) are compared. Scale bar=10 μm. In ii, the change is quantified using two-way ANOVA and a type III model fit (Armitage 2001). There was a statistically significant interaction between age and mutation status (F (1, 28)=12.89 *P*=.0012), no significant influence of age (F (1, 28)=180.76, *P*<.392) and no significant influence of mutation status (F (1, 28)=2.88, P=.1006). *P*-value shown on the graph stems from a *post-hoc* Tukey multiple comparison test. 4 mpf: n_+/+_=6, n_−/−_=6, 8 mpf: n_+/+_=5, n_−/−_=5, 13 mpf: n_+/+_=6, n_−/−_=4 Bars represent mean +/− SEM.

### Rad51-/- fish show a hyperproliferative phenotype in the kidney marrow

The number of cells in the WKM is determined by the balance between their proliferation and apoptosis rate, as well as their migration to the PB. To test whether the decrease in kidney cell numbers in *rad51* mutants was due to apoptosis, we carried out an AnnexinV-Propodium Iodide (AV-PI) staining assay on kidney tissue (Figure 3A and Figure S4A and B). We observed no statistically significant difference between *rad51*+/+ and -/- fish, excluding apoptosis as an initial cause for the decrease in cell numbers in the kidney. Therefore, we next focused on the proliferation of the kidney marrow cells.

**Figure 3.**
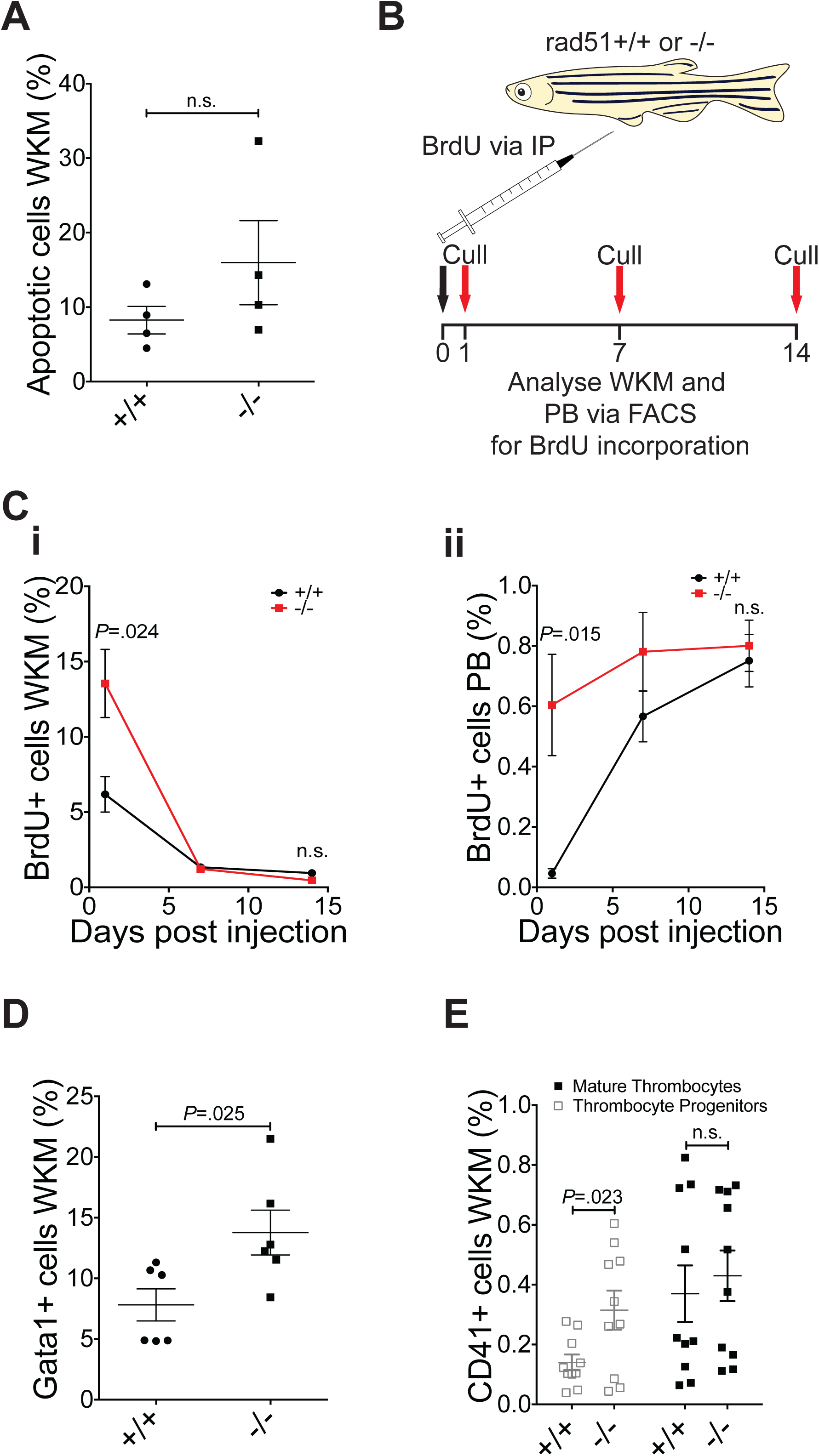
HSPCs in the kidney adult rad51 mutant fish show increased proliferation. (A) AnnexinV-PI assay to assess apoptosis in the kidney. Two-sided T-test, n_+/+_=4, n_−/−_=4. (B) Schematic of the experimental design for the BrdU incorporation experiments. Fish were injected once with 10 mg/ml BrdU and culled after 1, 7 or 14 days to obtain the blood and kidney marrow for antibody staining and FACS analysis. (C) Percentage of BrdU+ cells in the kidney (i). Two sided Student’s t-test, *P*=.024 at 1 dpi and *P*>.05 at 14 dpi. 1 dpi n_+/+_=5, n_−/−_=6; 7 dpi n_+/+_=6, n_−/−_=6; 14 dpi n_+/+_=5, n_−/−_=5. Percentage of BrdU+ cells in the peripheral blood (ii). Two sided Student’s t-test, *P*=.0015 at 1 dpi and *P*>.05 at 14 dpi. 1 dpi n_+/+_=5, n_−/−_=6; 7 dpi n_+/+_=6, n_−/−_=5; 14 dpi n_+/+_=5, n_−/−_=5. (D) Percentage of gata1:GFP+ cells in the kidney at 4 mpf. Two sided Student’s t-test, *P*=.025, n_+/+_=6, n_−/−_=6. (E) Percentage of dim and bright cd41:GFP+ cells in the kidney at 4 mpf, labeling thrombocytic progenitors and mature thrombocytes respectively. Two-tailed Student’s t-test. Thrombocytic progenitors: *P*=.023, mature thrombocytes: *P*=n.s., n_+/+_=10, n_−/−_=10. Bars represent mean +/- SEM in all graphs.

The standard assay for assessing cell proliferation rates is the incorporation of BrdU into the DNA. To follow the kinetics of BrdU incorporation and dilution via division and/or migration to the circulation, we measured BrdU labeling at several time points post injection (Figure 3B). Since erythrocytes are nucleated in zebrafish, our analysis was not limited only to leukocytes and allowed us to robustly assess changes in WKM as well as in PB. The initial number of BrdU+ cells in the WKM was two-fold higher in rad51-/- fish compared to the control (Figure 3Ci and Figure S4F), suggesting an increased proliferation rate in the mutant. This was followed by a fast dilution of the BrdU label over two weeks, because of the increased cell division of mutant HSPCs (Figure 3Ci). In line with this, the initial number of BrdU+ cells in the PB was higher in *rad51*-/- fish compared to wt siblings (Figure 3Cii), but then plateaued faster due the dilution of the label in the kidney (Figure 3Cii).

To show that blood, rather than other cells in WKM are proliferating, we generated mutants in various transgenic backgrounds. The rad51-/- *Tg(gata1a:GFP)* line(39), was used to assess the erythrocytic lineage, including erythrocytic progenitors. The *rad51*-/- *Tg(cd41Atga2b:EGFP)(40)* line was employed to detect thrombocytic progenitors, which are labeled in the *cd41CD41*:*EGFP^dim^* subpopulation(41). Consistent with the BrdU incorporation experiments, we saw an increase in newly made *gata1*:GFP+ erythrocytes in the kidney (Figure 3D and Figure S4C), as well as an increase in *cd41*:*EGFP*^dim^ thrombocytic progenitors (Figure 3E and Figure S4D).

Together, this evidence shows that loss of *rad51* leads to a hyperproliferation of HSPCs, possibly as a compensatory mechanism to prevent cytopenia in the PB. However, the lowered cellularity of the kidney in adult *rad51*-/- fish cannot be explained by this finding, suggesting that the kidney cytopenia stems from an early, possibly embryonic HSPC defect.

### Lack of Rad51 causes an HSPC defect during early development

The definitive wave of hematopoiesis starts at around 30 hpf when long-term HSCs are formed from endothelial cells of the ventral wall of the dorsal aorta. These newly made HSCs move to the caudal hematopoietic tissue (CHT), which serves as an intermediate place of hematopoiesis in which the HSCs expand greatly, akin to the mammalian fetal liver(42).

To assess the underlying cause of the decreased number of cells in the adult kidney of FA fish, we focused on embryonic hematopoiesis. To this end, we utilized whole-mount *in-situ* hybridisation (ISH) using a *cmyb*-specific probe, which labels HSPCs. Indeed, at 2 dpf, *rad51*-/- embryos had a decreased number of HSPCs compared to the wt embryos from the same clutch (Figure 4Ai and ii). At 4 dpf, the difference in the number of HSPCs in the CHT of *rad51*-/- and *wt* embryos was further exacerbated (Figure 4Bi and ii).

**Figure 4.**
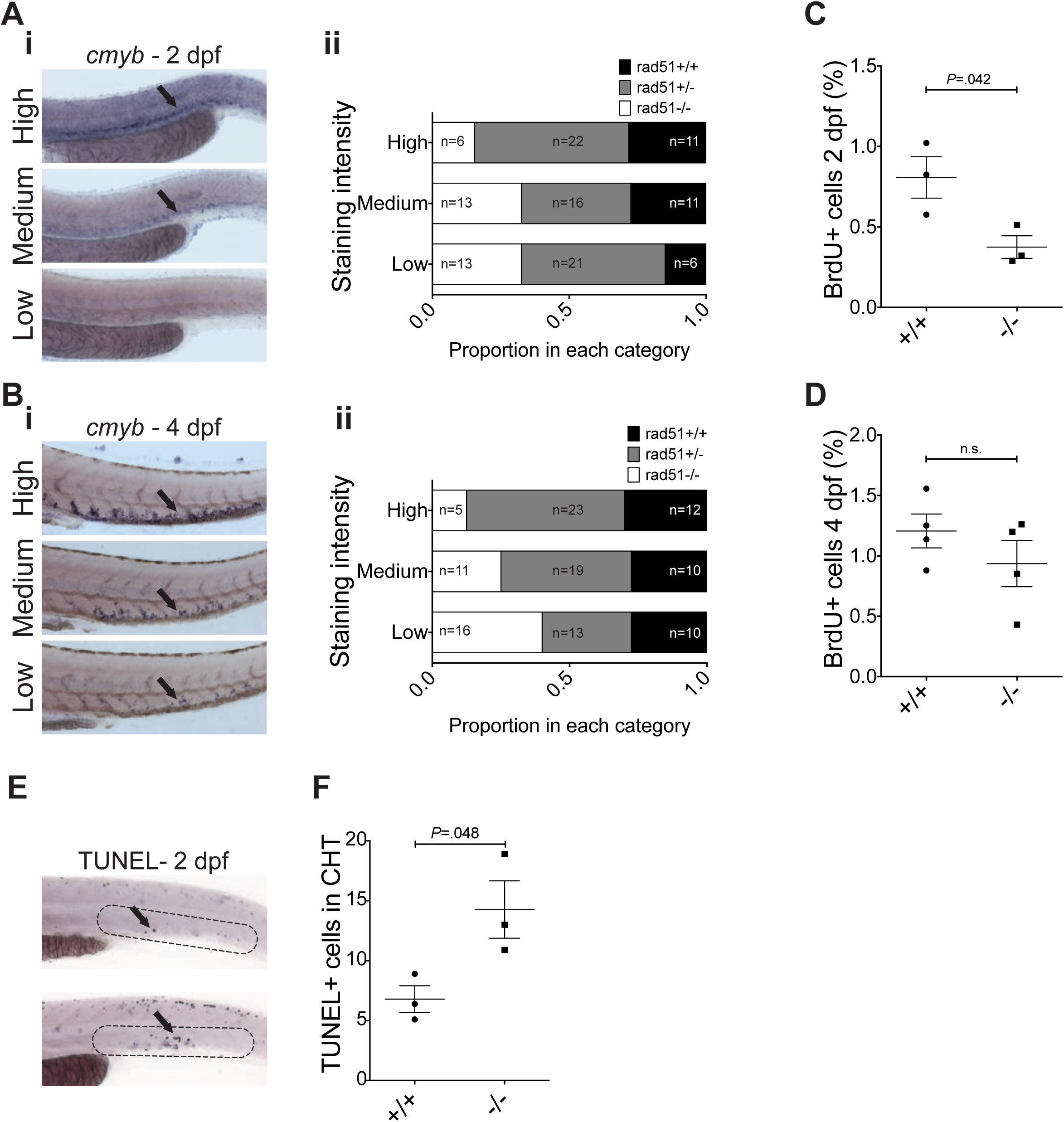
The *rad51^sa23805^* HSPC defect starts during embryonic development. (A) In-situ hybridization using a *cmyb*-specific probe at 2 dpf; The arrow shows HSPCs. Representative images of the three different staining categories are shown on the left and a quantification of the different genotypes on the right, n=119 from 2 clutches. (B) Quantification of BrdU+ cells in the tail at 2 dpf. Two-sided Student’s t-test, *P*=0.042, n_+/+_=3, n_−/−_=3. (C) In-situ hybridization using a *cmyb*-specific probe at 4 dpf; The arrow shows HSPCs. Representative images of the three different staining categories are shown on the left and a quantification of the different genotypes on the right, n=120 from 2 clutches. (D) Quantification of BrdU+ cells in the CHT at 4 dpf. Two-sided Student’s t-test, n_+/+_=4, n_−/−_=4. Bars represent mean +/− SEM in B and D. (E) Representative images of TUNEL-stained 2 dpf embryos from a *rad51*+/- incross. Dotted lines indicate the area of the CHT that was scored. Arrows indicate TUNEL+ cells. (F) Quantification of 3 clutches of TUNEL-stained 2 dpf *rad51*+/- incrosses. Each clutch was scored blindly and consisted of 10 +/+ and 10 −/− embryos each. Shown is the mean of all clutches +/− SEM.

Following up on that finding, we carried out a BrdU incorporation assay on the tail tissue of 2 dpf embryos, which showed that the proliferation rate in the tail of *rad51*-/- embryos was about half that of *rad51*+/+ embryos (Figure 4C). Although not statistically significant this trend was still apparent in the CHT at 4 dpf (Figure 4D).

In addition to proliferation, we also investigated apoptosis at 2 dpf by carrying out a TUNEL assay on several crosses of *rad51* heterozygotes. This revealed a twofold increase in apoptosis in the CHT in mutants compared to wt embryos (Figure 4E and F). Taken together, our data imply that the cytopenia in the adult kidney is caused by an increase apoptosis of HSPCs as well as reduced proliferation during early embryogenesis, mainly before 4 dpf.

### The HSPC defects in *rad51*-/- fish are mediated via *p53*

The defects seen in FA have recently been linked to an aggravated *p53* response (Ceccaldi et al 2012). To focus on the role of *p53* in the HSPC defect we generated a zebrafish line carrying mutations in both *rad51* and *p53*. This double mutation was able to rescue the number of HSPCs in the CHT of 4 dpf embryos (Figure 5A and Table S3). Importantly, the number of cells in the adult WKM of *p53-/- rad51-/-* fish reached wt levels by 4 mpf (Figure 5B). Along with the rescued marrow cellularity, there was no difference in the proliferation of cells in the WKM of wt and double mutant fish at 4 mpf, as shown by a BrdU incorporation assay (Figure 5C).

**Figure 5.**
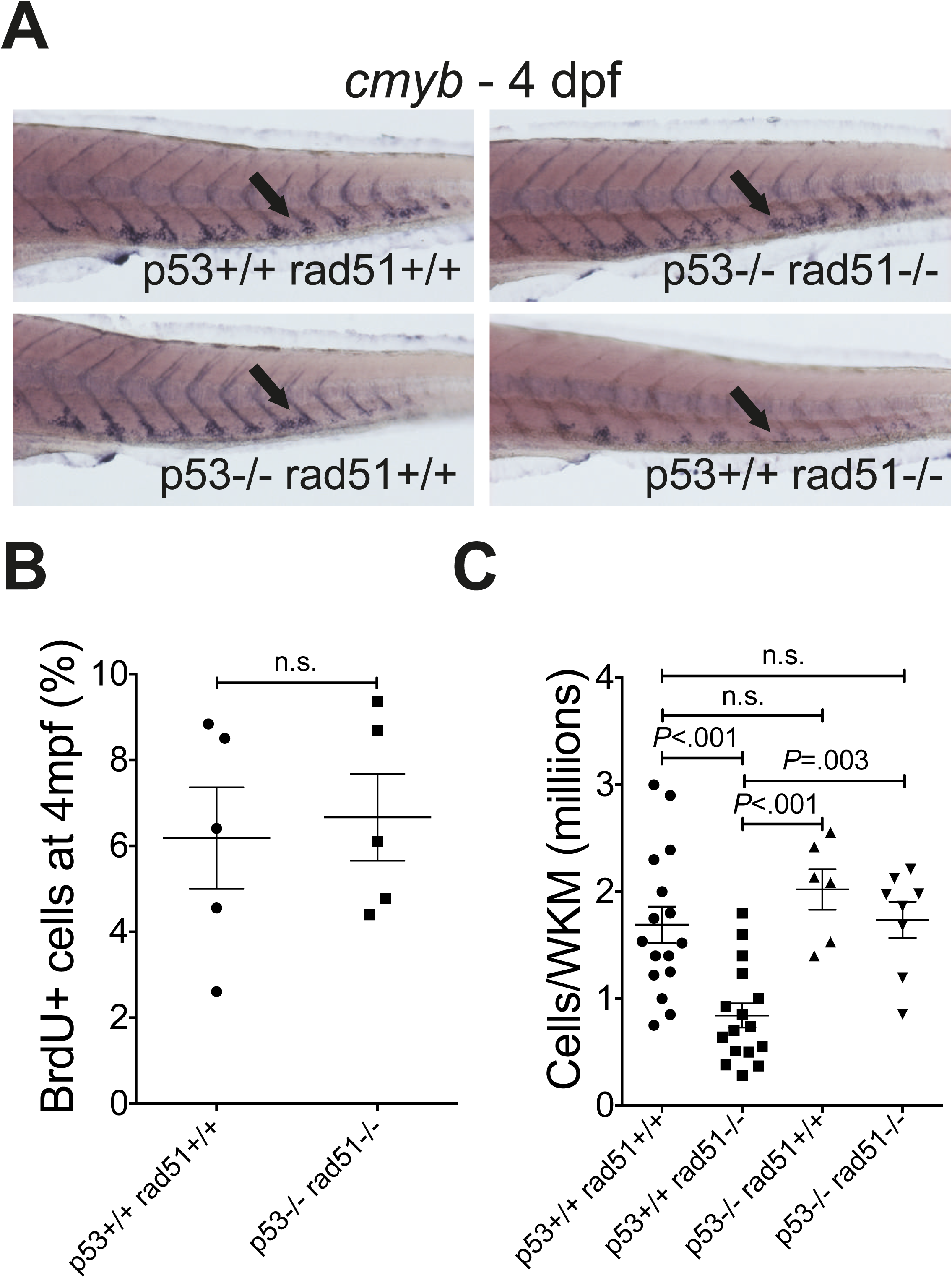
The HSPC defects in *rad51^sa23805^* fish are rescued in a *p53* mutant background. (A) Representative images of 4 dpf embryos resulting from in-crosses of *p53*+/- *rad51*+/- parents stained using a *cmyb*-specific probe. The total number of embryos used (all genotypes) n=237 from 4 clutches. For information about all genotypes please see Table S3. (B) Percentage of BrdU+ cells in the kidney at 4 mpf at 1 dpi. Two-sided Student’s t-test, n_*p53*+/+ *rad51*+/+_=5, n_*p53* −/− *rad51*−/−_=5. (C) Number of total cells per kidney at 4 mpf quantified using a hemocytometer. Analysis using one-way ANOVA (F (3,43=10.45), *P*<.0001), individual p-values shown in the figure are from Tukey’s *post-hoc* test, n_*p53*+/+ *rad51*+/+_=16,n_*p53*+/+ *rad51*−/−_=16, n_*p53*−/− *rad51*+/+_=6, n_*p53*−/− *rad51*−/−_=8. Bars represent mean +/− SEM in B and C.

We also examined the congenital phenotypes in the double mutants. The sex reversal seen in *rad51* single mutants was corrected (Figure S5A-C), but the size defect was not (Figure S5D). However, neither female, nor male double mutants were fertile (Figure S5A-C). Taken together, our data suggest that the marrow hypocellularity in adult fish was not due to the smaller size of *rad51* mutant, meaning that these two phenotypes are uncoupled. Instead, we observed a high correlation between the number of HSPCs generated early during embryonic development and the kidney marrow cellularity in adulthood. Therefore, the rescue in the embryonic definitive hematopoiesis can revert all defects observed in adult hematopoiesis in *rad51* mutants.

Finally, the tumor incidence of the double mutants was 30%, with the first tumors developing from 5 mpf (five months earlier than reported for *p53* single mutants(43)). The tumors resembled malignant peripheral nerve sheath tumors (MPNSTs), (Figure S5E-F), which is the most common type of malignancy in *p53*-/- fish (43). None of the other fish (i.e. wt, *rad51* mutants and heterozygotes) developed any kind of noticeable tumor, including the oldest 13 mpf *rad51*-/- fish.

### Rad51-/- fish are more sensitive to inflammatory stress

An aberrant inflammatory response has been postulated to be one of the potential causes of the BMF in FA. This is thought to be due increased expression of inflammatory cytokines in FA patients and an excess apoptosis of HSPCs in response to these factors. To test this hypothesis, we developed a zebrafish model of prolonged inflammatory stress. Over a period of four weeks, we injected rad51-/- and wt fish with the immunostimulant polyinosinic:polycytidylic acid (pI:pC) and assessed changes in the kidney cellularity, lineage output and the expression of genes associated with inflammation (Figure 6A).

**Figure 6.**
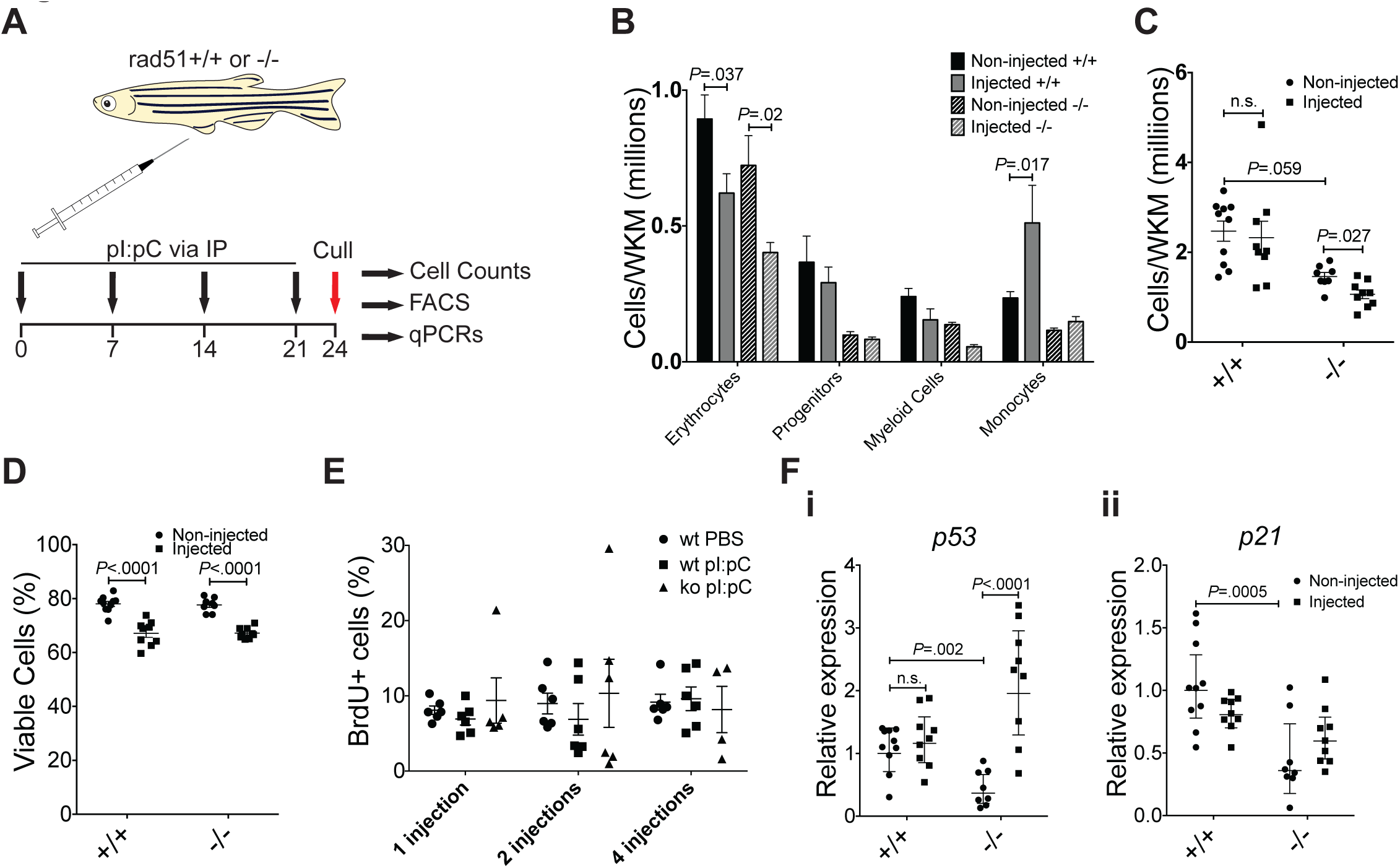
Lack of Rad51 causes increased sensitivity to prolonged inflammatory stress. (A) Schematic of the experimental design. Both wild type and *rad51*-/- fish were injected every seven days with 10 μ! 10 mg/ml polyinosinic:polycytidylic acid, four injections in total. All fish were culled 3 days after the last injection. Control fish were not injected; Rad51+/+, n_non-injected_=10, n_injected_=9; Rad51-/-, n_non-injected_=8, n_injected_=9. (B) Absolute number of cells belonging to different blood lineages in the kidney gained by combining FACS data with the cell counts shown in A. Statistical tests were carried out individually for each cell type, using two-way ANOVA. *P*-value shown on the graph stems from a *post-hoc* Šidak multiple comparison test, comparing non-injected to injected fish within each genotype. For all groups, n is the same as in A. (C) The total number of cells in the kidney in injected and non-injected fish. Two-way ANOVA was carried out on the reciprocal of the data to fulfill the requirement of homoscedasticity as measured by Bartlett’s test (before transformation: *P*=.0002, after transformation: *P*=.095). There was a statistically significant effect of mutation status (F (1,32)=29.86, *P*<.0001) and of injection status (F (1,32)=6.778, *P*=.014). P-value shown on the graph stems from a *post-hoc* Tukey multiple comparison test. For all groups, n is the same as in A. (D) Viable cells as determined by PI-staining. Bars represent mean +/− SEM in B, C and D. (E) Percentage of BrdU+ cells in the WKM of wild type (wt) and mutant (ko) fish in response to pI:pC. 1|=one injection, 2|=two injections, 4|=four injections. (F) Relative expression of genes linked to apoptosis and proliferation. (i) *p53*. *P*-value shown on the graph stems from a *post-hoc* Tukey multiple comparison test. (ii) *p21*. Bars represent geometric mean +/− 95% CI in E and F.

PI:pC resembles double-stranded RNA and is known to induce the expression of proinflammatory cytokines such as TNFα(44) and IL1 (45). Therefore, it has been considered to accurately mimic viral infection(45) in murine and fish models(44–47). Indeed, a single IP injection of pI:pC in zebrafish induced robust inflammatory response just six hours postinjection (Figure S6A). Mutants did not express more inflammatory cytokines when unchallenged (Figure S6B). Importantly, the prolonged exposure of adult wt fish to pI:pC resulted in a twofold increased production of monocytes (Figure 6B) but did not overtly affected kidney marrow cellularity or cell viability (Figure 6C and D). In addition, we observed clear upregulation of monocyte specific genes (Figure S6C). The skew towards monocyte production in turn decreased the erythrocyte output and to a lesser extent other lineages. In contrast, repeated pI:pC injections of *rad51*-/- fish led to no significant change in the number of produced monocytes (Figure 6B) and a ~25% decrease in the total number of cells in the kidney (Figure 6C). As in the wt fish, the prolonged inflammation decreased the number of erythrocytes in the kidney marrow of *rad51*-/- fish thus possibly contributing to the overall reduction in kidney cellularity. This was mediated by a normalization of proliferation rates in the kidneys of pI:pC injected mutant fish (Figure 6E and S6D). Therefore, the prolonged inflammatory stress can lead to severe marrow defects in our *rad51*-/- zebrafish FA model not only in terms of the lineage output but also by affecting the total marrow cellularity.

Interestingly, the unchallenged *rad51* mutants downregulated *p53*, but showed an exaggerated reaction in *p53* expression to repeated inflammatory stress (Figure 6Fi). Fitting with this observation, *p21* expression followed a similar trend in gene expression in non-injected and pI:pC injected mutants (Figure 6Fii).

## Discussion

The Fanconi anemia genes encode proteins that function cooperatively in the Fanconi DNA repair pathway. Here we characterized the first viable vertebrate *rad51* loss of function mutant. Loss of *rad51* in zebrafish recapitulated many congenital features of FA, such as short stature and microphthalmia(1); as well as hematological defects, including marrow cytopenia and accumulation of macrocytic erythrocytes in circulation(1, 8). Most importantly, *rad51*-/- fish showed increased sensitivity to crosslinking agents, an absolute diagnostic criterion of FA(8).

Although loss of *Rad51* leads to an early embryonic death in mice(34, 35), zebrafish lacking *rad51* survive to adulthood. This is not entirely surprising, as fish lacking *brca2* also survive to adulthood(48, 49), whereas *Brca2* mutant mice die before birth(50, 51). It has been hypothesized that maternal mRNA contributes to embryonic viability of zebrafish mutants(48). Interestingly, *rad51*-/- fish, like *brca2*-/- fish show sex reversal that can be rescued upon *p53* co-mutation(48, 49). Additional *p53* loss further caused development of MPNSTs in both *brca2* and *rad51* mutants, which occurred considerably earlier and at the higher incidence than in *p53* single mutants(43, 48, 49, 52).

FA is genetically and phenotypically heterogeneous disorder, but bone marrow failure is the most common cause of death(1) with patients having lowered CD34+ progenitor cell numbers from birth(11, 53). This led to hypothesis that hematological defects in FA originate from an impairment of HSPCs during embryonic development, which leads to a decreased number of HSPCs at birth(11, 12). Here we provide the first *in-vivo* evidence that the decrease in HSPC numbers in adult fish indeed stems from a combination of decreased proliferation and increased apoptosis during embryonic development. This defect appears to be mediated via *p53(11)*, as our *p53*/*rad51* double mutants did not display any observable hematological defects in embryos or adults.

In agreement with our study, knockdown of *fancd2* in zebrafish embryos causes massive apoptosis in the whole body, associated with upregulation of genes in the *p53* pathway, as well as decreased expression of cyclins. Strikingly, co-knockdown of *p53* rescued both the cyclin down-regulation, as well as apoptosis(54), resembling the situation in our fish and providing further evidence for the importance of *p53* signalling. This phenotype is, however, only partly consistent with what was observed in murine fetal *Fancd2*-/- cells. In the murine model the reduced number of HSPCs was set off by p38 mediated reduced proliferation, without any involvement of apoptosis or p53 signaling(55). More research into the causes of the early HSPC defects and how they can be mitigated is warranted. Due to the external development and their transparency zebrafish embryos would be an ideal tool to discover compounds that can alleviate these defects.

The decreased WKM cellularity in *rad51* zebrafish mutants also mirrors defects seen in *Lig4* (a protein involved in non-homologous end-joining) mutant mice, which show decreased BM cell numbers, coupled to an approximately twofold increase in proliferation of LT-HSCs(56). This underlines the importance of repairing double stranded breaks in HSC maintenance and suggests hyperproliferation of blood progenitors might be a common mechanism to cope with decreased cell numbers in the kidney/bone marrow. However, like most murine FA models(57, 58), the fish never progressed to pancytopenia or spontaneous BMF/KMF, which is probably due to species differences in the HSC compartment, lifespan and rearing conditions. Nevertheless, together with the clinical data(30, 31), these facts provide further evidence for the designation of *RAD51* as *FANCR* and show that our *rad51*-/- fish are a suitable model for FA.

The progressive decline of HSC numbers in FA patients leads to BMF and there is a considerable evidence for an overproduction of inflammatory cytokines in patients and mouse models(27, 59–63). Furthermore, inflammatory stress (for example by repeated pI:pC injections) can induce BMF in mouse models of FA(64, 65). We did not observe an increase in inflammatory cytokines in our unchallenged *rad51* mutants, ruling out cytokine overproduction as a cause for hematological defects. We did, however, observe a 25% decrease in WKM cellularity after repeated of pI:pC injections, possibly due to decreased proliferation compared to unchallenged fish. Unlike wt fish, *rad51* mutants were unable to respond appropriately to inflammation and increase monocyte production. The decrease in WKM cellularity as a result of inflammatory stress was accompanied by an upregulation of *p53*, again underlining the high importance of this pathway in the etiology of FA and explaining the normalized proliferation.

Our study characterized the first viable vertebrate model of *RAD51* loss, which recapitulates many human FA symptoms and is thus also the first zebrafish model of the disease. Further study of this mutant will increase our knowledge of the cellular roles of RAD51 *in-vivo* and will deepen our understanding of the molecular pathology of FA. Finally, the suitability of zebrafish embryos for high-throughput screening will significantly impact the development of novel therapeutics to improve HSPC function in FA patients.

## Methods

### Zebrafish care and strains

Fish lines were maintained in the Sanger Institute zebrafish facility according to EU regulations. Wild type fish were of the Tübingen long fin strain. Fish were genotyped as described previously(36).

### Western blotting

Western blotting was carried out as described previously(66). Antibodies can be found in supplementary Table 1.

### Chromosome spreads

Embryos were treated with 1 μg/ml mitomycin C (Sigma Aldrich) in egg water between 4 and 24 hpf. From here on we followed the Zebrafish Book 4^th^ edition(67), but kept the head of each embryo for genotyping. VECTASHIELD mounting medium with DAPI was used to visualize the spreads.

### Embryo irradiation

Embryos were irradiated at 24 hpf in a Gammacell 1000 Elite Blood Irradiator (MDS Nordiron) at 750 cGy.

### Immunostaining

Staining was carried out as described(68). We used Hoechst 33342 as nuclear stain. Embryos were imaged on a Leica SP-5 confocal microscope using a 40X water immersion lens. Antibodies can be found in Table S1.

### Histology

Formalin fixed tissues were processed and sectioned using standard techniques(69) followed by staining with Harris hematoxylin and eosin (H&E).

### AnnexinV-PI assay

We used the Alexa Fluor^®^ 488 Annexin V/Dead Cell Apoptosis Kit (Thermo Fisher) according to the manufacturer’s instructions.

### BrdU assay on adults

Fish were injected with 10 μl of 10 mg/ml 5-bromo-2∲-deoxyuridine (BrdU, Sigma Aldrich) and culled at 1, 7 or 14 days post injection. We extracted kidney and blood, made single cell suspensions and fixed cells in 70% EtOH overnight. BrdU immunostaining was carried out as described previously(66), after which the cells were resuspended in PBS and analyzed using FACS.

### BrdU assay on embryos

Embryos were chilled on ice for 15 minutes. This was followed by a 20-minute incubation in 10 mM BrdU on ice. After a 3 h recovery in egg water, embryos were fixed in 4% PFA. Heads from embryos were used for genotyping. For 2 dpf embryos, the whole tails were pooled according to genotype, whereas for 4 dpf embryos just the CHT was dissected. Samples were treated with 10mM DTT in 1X Danieau’s solution for 30 minutes at room temperature, followed by incubation in liberase (Roche) in PBS for 3 hours at 37 °C. The reaction was stopped by replacing the solution with 5%FBS/PBS. Single cell suspensions were fixed in 70% ethanol overnight. From here on, the staining process and analysis was identical to cells obtained from adults. Antibodies can be found in Table S1.

### TUNEL assays

Embryos for TUNEL assays were fixed and stored as for ISH. Staining was carried out using the In Situ Cell Death Detection Kit, AP (Roche) according to the manufacturer’s instructions.

### Kidney FACS

Dissected kidneys were placed in 5%FBS/PBS and processed to single cell suspensions. Dead cells were excluded using 4∲,6-diamidino-2-phenylindole (DAPI, Sigma-Aldrich Inc.) or propidium iodide (PI). Flow cytometry was carried out on a MoFlo XDP (Beckman Coulter), a BD LSRFortessa, or a BD Influx (BD Biosciences).

### *In-situ* hybridization (ISH)

In-situ hybridization was carried out as described previously(70). We used a *cmyb* antisense probe. Embryos were blindly sorted into high, medium and low staining conditions followed by genotyping.

### Long-term pI:pC injections

Fish were injected with 10 mg/ml polyinosinic:polycytidylic acid (pI:pC, Sigma-Aldrich) once a week, totaling 4 injections. Fish were culled one or three days after the last injection. Optionally, 10 mg/ml BrdU were added to the last injection. Kidneys were processed to single cell suspensions for FACS, as described above and cell numbers counted. Remaining cell suspensions were used for gene expression analysis.

### qPCRs

The qPCR reaction used SYBR green (Thermo Fisher) and run on a QuantStudio 3 (Thermo Fisher) qPCR machine. Primers used are listed in Table S2.

## Acknowledgements

We thank the Sanger Institute Zebrafish Mutation Project for supplying the *rad51^sa23805^* allele. We would like to thank Sebastian Gerety for supplying the *tp53^zdf1^* line, Yvette Hooks for her help with histology, the Sanger Institute FACS core facility, as well as Debbie Goode and Charlotte Labalette for their experimental help.

This work was supported by Cancer Research UK Grant C45041/A14953 (to A.C.), a core support grant from the Wellcome Trust and Medical Research Council to the Wellcome Trust-Medical Research Council Cambridge Stem Cell Institute, as well as a EHA-Jose Carreras Foundation Young Investigator Award and Isaac Newton Trust grant (to A.C.).

## Authorship Contribution

J.G.B., E.B.M. and L.F. carried out experiments. J.G.B., E.B.M. and A.C. analyzed data. J.G.B and A.C. designed figures and wrote the manuscript.

